# Characterization Of Ancestral Origin Of Cystic Fibrosis Of Patients With New Reported Mutations In CFTR

**DOI:** 10.1101/2020.05.06.081653

**Authors:** César Paz-y-Miño, Ana Karina Zambrano, Juan Carlos Ruiz-Cabezas, Isaac Armendáriz-Castillo, Jennyfer M. García-Cárdenas, Santiago Guerrero, Andrés López-Cortés, Andy Pérez-Villa, Patricia Guevara-Ramírez, Verónica Yumiceba, Paola E. Leone

**Affiliations:** Centro de Investigación Genética y Genómica, Facultad de Ciencias de la Salud Eugenio Espejo, Universidad UTE, Av. Mariscal Sucre and Mariana de Jesús, Block I, Quito 170129, Ecuador; Universidad de Especialidades Espíritu Santo (UEES). Guayaquil-Ecuador; Instituto de Biomedicina, Universidad Católica de Santiago de Guayaquil. Guayaquil – Ecuador; Instituto Oncológico Nacional de la Sociedad de Lucha Contra el Cáncer (ION-SOLCA). Guayaquil-Ecuador

**Keywords:** Cystic Fibrosis, Native American, Ecuadorian, CFTR

## Abstract

The incidence of Cystic fibrosis (CF) and the frequency of the variants for CFTR depend on the population; furthermore, CF symptomatology is characterized by obstructive lung disease, pancreatic insufficiency among others, reliant on the individual genotype. Ecuadorian population is a mixture of Native Americans, Europeans, and Africans. That population admixture could be the reason for the new mutations reported in a previous study by Ruiz et al. (2019). A panel of 46 Ancestry Informative Markers was used to estimate the ancestral proportions of each available sample (12 samples in total). As a result, the Native American ancestry proportion was the most prevalent in almost all individuals, except for three patients from Guayaquil with the mutation *[c.757G>A:p.Gly253Arg; c.1352G>T:p.Gly451Val]* who had the highest European composition.

## Introduction

Cystic fibrosis (CF) is an autosomal recessive disorder that has been extensively studied among populations (1). It is characterized by obstructive lung disease, pancreatic insufficiency, diabetes, liver disease, among others (2). The most frequent worldwide mutation in Cystic Fibrosis Transmembrane Conductance Regulator (CFTR) protein gene is c.1521_1523delCTT (p.Phe508del) (3) which originated between 11 000 and 34 000 years ago in Europeans, then it spread across all Europe (4). CF occurs in 1 out of 2 500 live births with high prevalence in the European ancestry, and the frequency of the heterozygotes has been reported as 1 in 25 in Europeans (5) (6). There are plenty of studies in CF, yet the majority in Europeans, underrepresenting the Latin Americans (4) (6) (7) (8). In the United States, a study reported the CF incidence to be 1 in 9 200 Hispanics and 1 in 10 900 Native Americans, yet the USA has a different population structure to South America(1) (6) (9). In general, in Latin America the incidence is 1 per 6 000 newborn alive; specifically, Ecuador exhibits an incidence of 1 in 11 252 newborns (10) (11) (12).

Ecuadorian population, located in the Northwest of South America is a mixed population conformed by Native Americans, Europeans who arrived in the 16^th^ century during the conquest, and Africans who came with them as slaves. According to the last census, the population projection for 2020 was estimated as 17 510 643 Ecuadorians. Moreover, Ecuadorian self–identified as: “mestizos” 71.9%, “montubios” 7.4%, Afro-Ecuadorian 7.2%, “indígenas” 7%, “blancos” 6.1% and others (0.4%) (13). There are also reports of the Ecuadorian ancestry using AIMs in mestizo population where Native American was the most prevalent ancestry (59.6%), followed by European (28.8%) and lastly African (11.6%) (14) (15).

Like other South American studies, Ecuador is underrepresented in cystic fibrosis research, and none of them involve the comparison of the mutations with the population’s origin. Paz-y-Miño et.al (1999) reported 10 cases of Ecuadorian CF patients, at least 60% of the mutations differ from c.1521_1523delCTT (p.Phe508del) (16). Valle et al. (2007), analyzed 62 Ecuadorian CF patients, the most prevalent mutation was F508del (37.1%) (12). The last report by Ortiz et al. (2017), included 48 Ecuadorian individuals with CF, reported F508del with the highest frequency (20.27%)(17). These studies, however, are mainly focused on the particular F508 mutation, revealing that the percentage is not relatively high as in Europeans. The incidence and the frequency of the CF mutation depend on the population under study, Ecuadorians are a mestizo population, and the population’s composition is not clear yet.

Here we provide the ancestry origin data of 46 Ancestry Informative Markers of the individuals with the new mutations reported in a previous study of CF patients from Ecuador (18). We aimed to elucidate if the mutations reported are mainly from European ancestry, due to the previous data of the main incidence.

## Methods

### Samples and DNA extraction

Twelve CF patients from Guayaquil (coast) and Cuenca (highland) who were available and had new CFTR disease-causing variants reported in a previous study were selected: one patient from Guayaquil with *c.1473T>A:p.Cys491*;* one patient from Guayaquil and two from Cuenca with *c.2672del:p.Asp891Alafs*15*; one patient from Cuenca with *c.1486T>C:p.Trp496Arg*; six patients from Guayaquil and one from Cuenca with [*c.757G>A:p.Gly253Arg; c.1352G>T:p.Gly451Val]*, were selected (18). DNA was extracted using Chelex 100 (Bio-Rad) (10%) from peripheral blood samples collected on FTA cards (GE Healthcare Life Sciences) and quantified using NanoDrop (Thermo Scientific). To protect the identity of the individuals, the samples were anonymized.

### DNA amplification

PCR amplification of the twelve CF samples and controls (positive: 2800 and negative) was performed using 46 AIMs-INDELs: MID-1470, MID-777, MID-196, MID-881, MID-3122, MID-548, MID-659, MID-2011, MID-2929, MID-593, MID-798, MID-1193, MID-1871, MID-17, MID-2538, MID-1644, MID-3854, MID-2275, MID-94, MID-3072, MID-772, MID-2313, MID-397, MID-1636, MID-51, MID-2431, MID-2264, MID-2256, MID-128, MID-15, MID-2241, MID-419, MID-943, MID-159, MID-2005, MID-250, MID-1802, MID-1607, MID-1734, MID-406, MID-1386, MID-1726, MID-3626, MID-360, MID-1603, MID-2719 (19), in one multiplex reaction and following the standardized protocol of the laboratory. The fragment separation was carried out in 3500 Genetic Analyzers (Applied Biosystems). Data were collected with Data collection v3 and visualized with Gene Mapper v5.

### Statistical Analyses

Data were analyzed with Structure v2.3.4 in order to estimate the ancestral proportions in the population, the runs consisted of a burn-in length of 10 000 followed by 10 000 Markov Chain Monte Carlo (MCMC) interactions. The option used was the admixture model (“Use population information to test for migrants”). The cluster considered for the analysis was one to three (k=1, k=2, and k=3) due to the historical background of the Ecuadorian population and according to the cluster identification by Evanno *et al* (20) (14).

Principal component analysis (PCA) was built with RStudio v1.1.453 to visualize the CF individuals’ structure: the correlation between individuals under analysis and the reference population from HGDP-CEPH (Native Americans, Europeans, and Africans) subset H952 (19).

## Results

The DNA quantification was optimal to perform the PCR (5-20ng/ul). After the amplification, complete profiles were obtained. A total of 339 individuals (reference population and samples), were analyzed, assuming a clustering of three using the information to test for migrants, permuting 10 000 burn-in periods and 10 000 interactions and a bar plot was obtained showing the main ancestral population analyzed (Fig 1).

**Fig 1.**
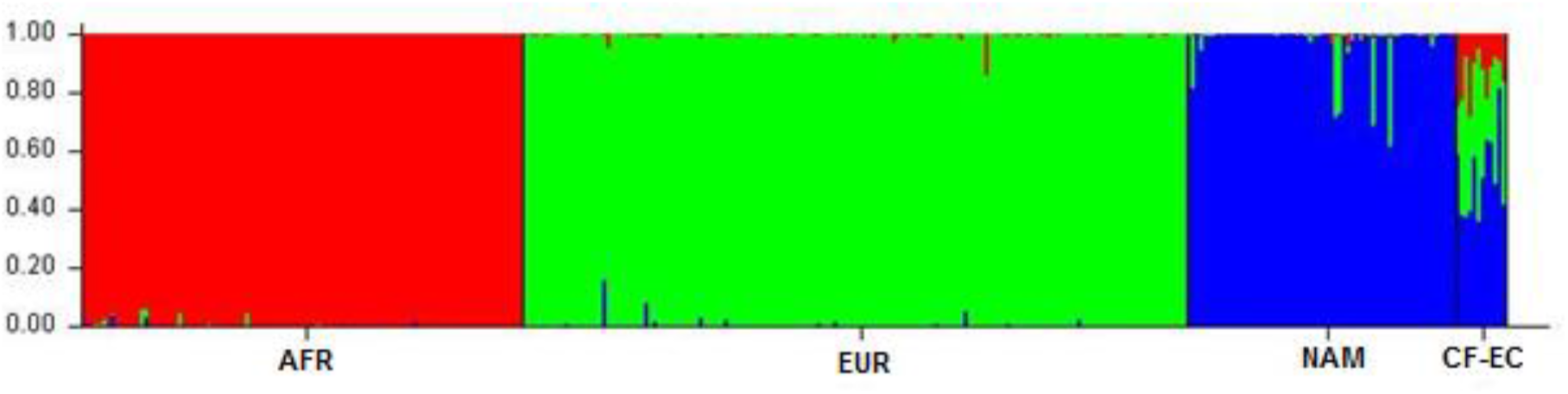
Bar plot grouped by population identification (AFR: African ancestry, EUR: European ancestry, NAM: Native American ancestry, CF-EC: Ecuadorian Cystic Fibrosis patients). Three inferred clusters (K=3).

Principal component analysis (PCA) results showed the three reference populations clearly differentiate between them. The CF Ecuadorian population is in the middle of them but mainly between European and Native American reference population. The two main principal components represented 38.86% of the total (Fig 2).

**Fig 2.**
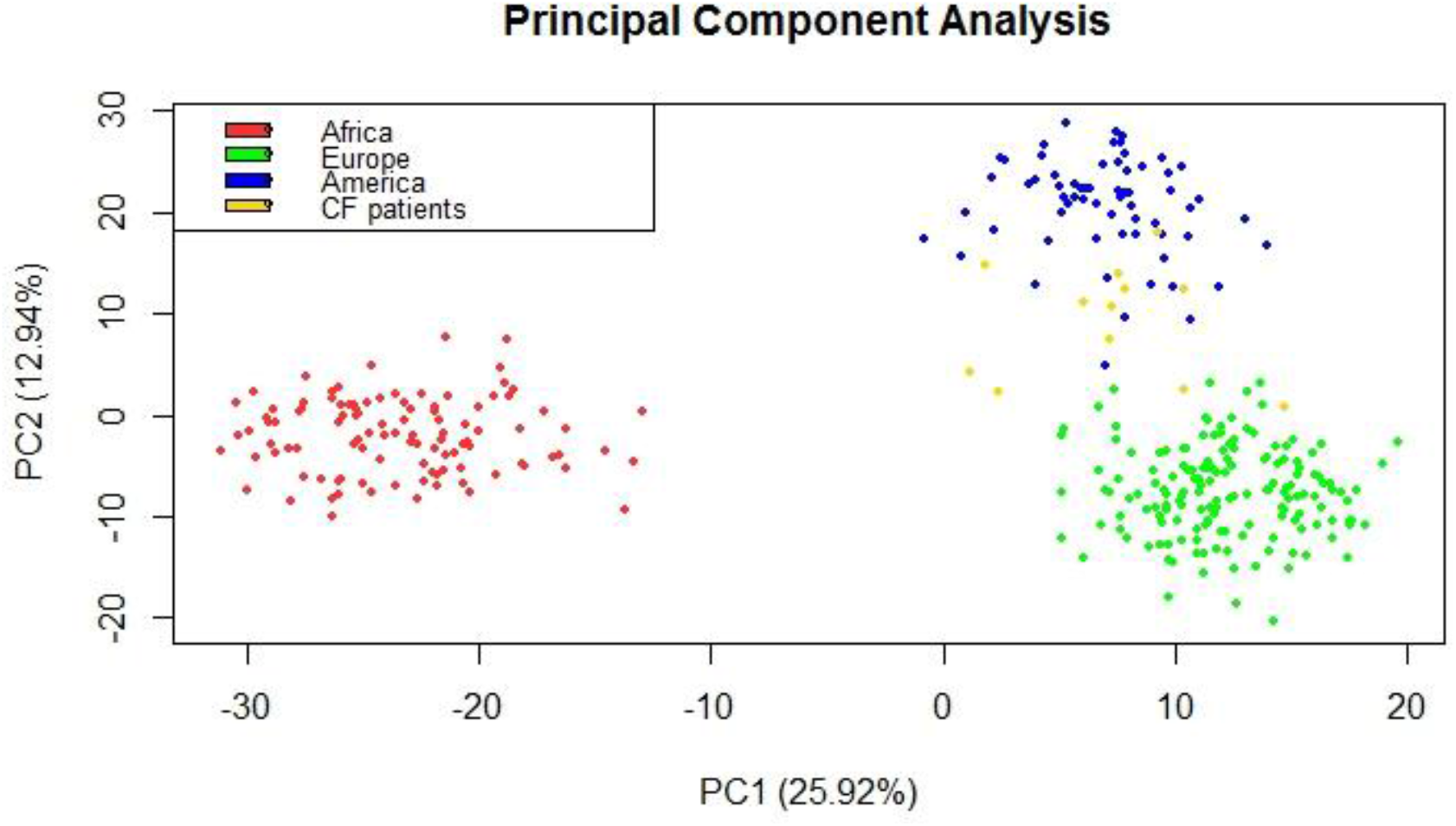
Principal component analysis of cystic fibrosis Ecuadorian patients.

A percentage of the ancestral composition of each individual was obtained, as a result, a heterogeneous percentage was found depending on the individual and the region under study. Hence, clearly showing the admixture of the Ecuadorian population according to history (Table 1). The global ancestry composition of CF patients was: The Native American 50% (standard deviation of 14.03), the European 35% (standard deviation of 15.5) and the African 11.5% (standard deviation of 7.82). The Native American ancestry was the first origin of almost all individuals, except for three patients from Guayaquil with the mutation *[c.757G>A:p.Gly253Arg; c.1352G>T:p.Gly451Val]* with the highest European composition.

**Table 1.** Percentage of the ancestral composition of each individual under study.

| PATIENT | AFR | EUR | NAM | MUTATION | MUTATIONS' REFERENCE |
| --- | --- | --- | --- | --- | --- |
| 01 | 24.6 | 16.7 | 58.7 | *c.1473T>A:p.Cys491\** | (18) |
| 02 | 7.6 | 43.5 | 48.9 | *c.2672del:p.Asp891Alafs\*15* | (18) |
| 03 | 21.6 | 14.8 | 63.6 | *c.2672del:p.Asp891Alafs\*15* | (18) |
| 04 | 16 | 42.2 | 41.8 | *c.2672del:p.Asp891Alafs\*15* | (18) |
| 05 | 9.1 | 8.9 | 82 | *c.1486T>C:p.Trp496Arg* | (18) |
| 06 | 22.5 | 39.6 | 37.9 | [*c.757G>A:p.Gly253Arg; c.1352G>T:p.Gly451Val*] | (18) |
| 07 | 7.2 | 55.3 | 37.5 | [c.757G>A:p.Gly253Arg;<br>c.1352G>T:p.Gly451Val] | (18) |
| 08 | 28.2 | 31.9 | 39.9 | [c.757G>A:p.Gly253Arg;<br>c.1352G>T:p.Gly451Val] | (18) |
| 09 | 5 | 58.7 | 36.3 | [c.757G>A:p.Gly253Arg;<br>c.1352G>T:p.Gly451Val] | (18) |
| 10 | 11.9 | 37.2 | 50.9 | [c.757G>A:p.Gly253Arg;<br>c.1352G>T:p.Gly451Val] | (18) |
| 11 | 9.3 | 32.1 | 58.6 | [c.757G>A:p.Gly253Arg;<br>c.1352G>T:p.Gly451Val] | (18) |
| 12 | 11 | 25.7 | 63.3 | [c.757G>A:p.Gly253Arg;<br>c.1352G>T:p.Gly451Val] | (18) |

\* AFR: African ancestry, EUR: European Ancestry and NAM: Native American Ancestry.

## Discussion

The present study is the first report of the ancestral composition of CF Ecuadorian patients with new CFTR mutations. There are plenty of CF studies among different populations that revealed the differences between gender, age, and symptoms in CF patients. Some studies compared CF patients of different ages and gender describing that the incidence of CF in Europe is higher in children than in adults, approximately 4 CF children per 3 adults and around 1.1 males per each female (21) (22).

There are reports that evidenced CF prevalence, differ depending on ethnicity. For instance, the incidence in Native Americans, whites and black individuals is 37.2, 38.8 and 17.1 per 100 000, respectively (23). Moreover, other studies revealed the incidence among diverse ethnicities: as an example, the prevalence of CF reported by Rohlfs et al. (2011) is 1 in 242 in Asian, 1 in 28 Caucasian, 1 in 59 Hispanic and 1 in 70 Native American (2). That study clearly revealed the ethnic differences in the incidence and the distribution of CF worldwide.

In Ecuador, there are studies about the ancestral origin, for instance, the Ecuadorian was reported to be composed of 59.6% of Native American, 28.8% of European and lastly 11.6% of African origin (14) (15).

In addition to the variable predisposition of CF among populations, reports exhibit a total of 2 063 mutations listed on the CFTR mutation database (24); while in the CFTR2 database, the most recent file updated on 8 December 2017, shows a total of 374 variants (25). Those variants were identified in different populations in diverse frequencies. For instance, the frequencies of the most common variant c.1521_1523delCTT (p.Phe508del) depend on ethnicity, it was reported as 72% in US Caucasians, ~41% in African Americans and 18% in Iranians, yet it also differs among Caucasians (26) (2) (27) (1) (28).

There are some mutations that have been commonly reported in ethnic groups, as examples, c.1624G>T (p.Gly542X) was reported in 43% of Turkish origin (29), while in a study in Peruvian patients the frequency was 6.9% (30); c.3846G>A (p.Trp1282Ter) was reported in 43% of Ashkenazi patients (31); c.2988+1G>A (3120+1G>A) reported in 12.3% of native African CF patients (32) (33); c.3909C>G (p.Asn1303Lys) described in 1.7% of the total number of CF analyzed from Europeans and the United States population (34), while in Algerian population the frequency was 20% (35); c.1652G>A (p.Gly551Asp) presented a frequency of 3% in North Brazilian population (36). Furthermore, some mutations have been found in a specific ethnic group, like, c.16C>G (p.Leu6Val) was found in one Argentinian, c.3294G>C (p.Trp1098Cys) was found in one Mexican, among other variations described (4); c.3276C>G (p.Tyr1092Ter) was found in Jews from Iraq (31) (2).

In conclusion, the identification of ethnicity-dependent mutations would be an important aspect of CF testing in Ecuador. The present study exhibited a greater ancestral composition of Native American, followed by European and lastly African, the mixed population origin could possibly explain the new CF mutations reported.

## Limitations

Although we have found the ancestral proportions of the majority CF patient with new mutations previously reported, we could not access all the samples due to the available conditions of the patients. Moreover, a larger CF patients study with the commonly reported mutation should be conducted to better approximation to the ancestral proportions of the patients.

## List of Abbreviations

CF: cystic fibrosis
CFTR: Cystic Fibrosis Transmembrane Conductance Regulator
AIMs: Ancestry Informative Markers
INDELs: Insertions Deletions
PCA: Principal Component Analysis
AFR: African ancestry
EUR: European ancestry
NAM: Native American ancestry

## Declarations

### Ethics approval and consent to participate

The patients were registered with the Ecuadorian Cystic Fibrosis Foundation at Guayaquil and Cuenca. Patients’ informed consent, including the paternal signed authorization to participate in the study. The investigation was approved with the number 2018-127E by “Comité de Ética de Investigación en Seres Humanos Universidad San Francisco de Quito”.

### Consent for publication

Not applicable.

### Availability of data

The data is presented in the manuscript.

### Authors’ contributions

César Paz-y-Miño: coordination and followed up with the development of the article.

Ana Karina Zambrano: design, experimental procedure, data analysis and writing.

Juan Carlos Ruiz-Cabezas: written edition.

Isaac Armendáriz-Castillo: written edition.

Jennyfer García-Cárdenas: written edition.

Santiago Guerrero: written edition.

Andrés López-Cortés: written edition.

Andy Pérez-Villa: writing and formatting

Patricia Guevara-Ramírez: written edition.

Verónica Yumiceba: written edition.

Paola E. Leone: written edition.

## Acknowledgments

The authors are grateful to the patients for their selfless participation in the present study. To all the people and institutions involved in the development of the present study.

